# G-quadruplexes are a source of vulnerability in *BRCA2* deficient granule cell progenitors and medulloblastoma

**DOI:** 10.1101/2024.07.20.604431

**Authors:** Danielle L. Keahi, Mathijs A. Sanders, Matthew R. Paul, Andrew L. H. Webster, Yin Fang, Tom F. Wiley, Samer Shalaby, Thomas S. Carroll, Settara C. Chandrasekharappa, Carolina Sandoval-Garcia, Margaret L. MacMillan, John E. Wagner, Mary E. Hatten, Agata Smogorzewska

## Abstract

Biallelic pathogenic variants in the essential DNA repair gene *BRCA2* causes Fanconi anemia, complementation group FA-D1. Patients in this group are highly prone to develop embryonal tumors, most commonly medulloblastoma arising from the cerebellar granule cell progenitors (GCPs). GCPs undergo high proliferation in the postnatal cerebellum under SHH activation, but the type of DNA lesions that require the function of the BRCA2 to prevent tumorigenesis remains unknown. To identify such lesions, we assessed both GCP neurodevelopment and tumor formation using a mouse model with deletion of exons three and four of *Brca2* in the central nervous system, coupled with global *Trp53* loss. *Brca2^Δex3-4^;Trp53^-/-^* animals developed SHH subgroup medulloblastomas with complete penetrance. Whole-genome sequencing of the tumors identified structural variants with breakpoints enriched in areas overlapping G-quadruplexes (G4s). *Brca2*-deficient GCPs exhibited decreased replication speed in the presence of the G4-stabilizer pyridostatin. *Pif1* helicase, which resolves G4s during replication, was highly upregulated in tumors, and *Pif1* knockout in primary MB tumor cells resulted in increased genome instability upon pyridostatin treatment. These data suggest that G4s may represent sites prone to replication stalling in highly proliferative GCPs and without BRCA2, G4s become a source of genome instability. Tumor cells upregulate G4-resolving helicases to facilitate rapid proliferation through G4s highlighting PIF1 helicase as a potential therapeutic target for treatment of *BRCA2*-deficient medulloblastomas.

## Introduction

Patients with biallelic pathogenic variants in the *BRCA2* gene belong to Fanconi anemia complementation group D1 (FA-D1) (Howlett et al. 2002; Wagner et al. 2004; Kennedy and D’Andrea 2005; Myers et al. 2012). Classically, these patients present with congenital abnormalities and are predisposed to leukemia and embryonal tumors at an early age, although presence of certain BRCA2 variants may result in milder disease (Howlett et al. 2002; Rickman et al. 2020; Radulovic et al. 2023). BRCA2 is a large 3418 amino acid protein that maintains genome stability through its dual roles in promoting homology-directed repair (HDR) of DNA double-strand breaks (DSBs) (Sharan et al. 1997; Powell et al. 2002; Prakash et al. 2015) and protecting stalled replication forks from excessive degradation by nucleases (Schlacher et al. 2011; Siaud et al. 2011; Schlacher et al. 2012).

BRCA2 has several interaction partners that facilitate its critical role in genome stability. BRCA2 interacts with the RAD51 recombinase through its BRC repeats and facilitates RAD51 loading and stable nucleoprotein filament formation on single stranded DNA (Yang et al. 2002; Yang et al. 2005; Jensen et al. 2010). The BRCA2 N-terminal domain contains an interaction site for PALB2 to allow for localization to DSBs and stalled forks and for efficient HDR (Xia et al. 2006; Murphy et al. 2014). Patients with biallelic PALB2 variants belong to the FA-N complementation group and have a predisposition to embryonal tumors of childhood, like FA-D1 patients (Offit et al. 2003; Reid et al. 2007; Xia et al. 2007).

Patients in the FA-D1 complementation group develop medulloblastoma (MB) as one of their earliest malignancies, and these tumors are typically classified as Sonic Hedgehog (SHH) subgroup (Offit et al. 2003; Hirsch et al. 2004; Alter et al. 2007; Myers et al. 2012; Miele et al. 2015; Waszak et al. 2018). MB arising from BRCA2-deficiency was previously modeled in mice with nervous system-wide *Brca2* mutation targeted to exon 11 that affected cerebellar development and led to medulloblastoma formation (Frappart et al. 2007; Frappart et al. 2009).

The cell of origin of SHH-MB is the granule cell progenitor (GCP) of the cerebellum (Salsano et al. 2004; Hovestadt et al. 2019; Vladoiu et al. 2019). These neuronal progenitors proliferate highly in the early postnatal period and give rise to the cerebellar granule cells, the most numerous neuron population in the brain (Azevedo FA 2009). GCPs undergo extensive clonal expansion in response to activation of the Sonic Hedgehog (SHH) signaling pathway in the early postnatal period (Gao et al. 1991; Wechsler-Reya and Scott 1999; Espinosa and Luo 2008). This is followed by cell cycle exit and migration from the external germinal layer (EGL) of the cerebellum to the inner granule layer (IGL) and differentiation into multipolar granule neurons (Consalez et al. 2020).

Constitutive SHH activation through loss of *PTCH1* in GCPs can lead to MB formation in mouse models through unchecked GCP proliferation (Oliver et al. 2005; Schuller et al. 2008; Yang et al. 2008). Analogously, germline pathogenic *PTCH1* variants associated with Gorlin syndrome lead to medulloblastoma in humans (Hahn et al. 1996). *In vitro*, activation of SHH signaling by treating GCPs with the N-terminal fragment of SHH (SHH-N) leads to faster DNA replication resulting in DNA damage in S-phase cells and increased origin licensing (Tamayo-Orrego et al. 2020). These findings suggest that GCPs may be especially sensitive to fork stalling at replication-blocking lesions during the proliferative phase of their development. Such stalling could be triggered by DNA lesions, proteins trapped on DNA, depletion of nucleotides required for ongoing DNA synthesis, and unresolved DNA secondary structures such as RNA/DNA hybrids and G-quadruplexes (G4s) that impede the replication machinery (Zeman and Cimprich 2014).

G4s are DNA secondary structures that distort the DNA helix and can stall replicative polymerases (Kaguni, 1982; Woodford, 1994; Weitzmann, 1996; Kamath-Loeb, 2001) and are linked to mutations in human cancers (Huppert 2008; De & Michor, 2011; Rodriguez, 2012). G4s may lead to increased DNA breakage in the absence of BRCA2 due to replication stalling (Xu et al. 2017; Groelly et al. 2022; Lee et al. 2022). G4s can be unwound by specific helicases including WRN (Fry and Loeb 1999; Kamath-Loeb et al. 2001; Crabbe et al. 2004), BLM (Sun et al. 1998; Johnson et al. 2010; Drosopoulos et al. 2015), FANCJ/BRIP1 (London et al. 2008; Castillo Bosch et al. 2014), and PIF1 (Ribeyre et al. 2009; Sanders 2010; Paeschke et al. 2011; Paeschke et al. 2013; Zhou et al. 2014). In this study, we highlight the intrinsic vulnerability of *Brca2^Δex3-4^* GCPs to replication stalling at G-quadruplex-rich loci that may lead to the mutations that drive medulloblastoma.

## Results

### Loss of the PALB2-binding domain of *Brca2* coupled with global *Trp53* loss leads to the development of SHH subgroup medulloblastoma

To identify endogenous lesions that may lead to MB development in patients with BRCA2 deficiency, we coupled the *Brca2*^Δ^*^ex3-4^* mouse model (Ludwig et al. 2001) with global *Trp53* knockout (Jacks et al. 1994). Conditional deletion of exons 3 and 4 of *Brca2* leads to breast carcinoma development when Cre expression is driven by the *Wap* promoter (Ludwig et al. 2001). It also results in replication stress, mitotic abnormalities, and G1 arrest in mammary epithelial cells (Feng and Jasin 2017). We expressed Cre under the control of human glial fibrillary acidic protein (*hGFAP)* promoter, leading to a conditional loss of exons 3 and 4 of *Brca2* in the developing mouse central nervous system (CNS) **(Figure 1A)**. We observed the loss of exons 3 and 4 using RNA sequencing and whole-genome sequencing of *Brca2*^Δ^*^ex3-4^* tumors and GCPs **(Figure 1B)**. However, the expression of BRCA2 protein appeared unchanged as assessed by western blotting using an antibody recognizing amino acids 1651-1812 **(Supplemental Figure 1A).** In this model, alternative splicing between exons 2 and 5 **(Figure 1B)** allows for expression of an alternate in-frame methionine in exon 7 leading to loss of 103 amino acids of the N terminus, a change that is not visible on a western blot **(Figure 1C).** Mutation of the BRCA2 N-terminus disrupts its PALB2-binding domain (Feng and Jasin 2017) and can lead to abrogated recruitment of BRCA2 to chromatin and diminished RAD51 filament formation (Xia et al. 2007). Indeed, *Brca2^WT^; Trp53^+/+^* GCPs but not *Brca2^Δex3-4^; Trp53^+/+^*GCPs induced RAD51 foci after treatment with mitomycin C (MMC) **(Figure 1D and E).**

**Figure 1.**
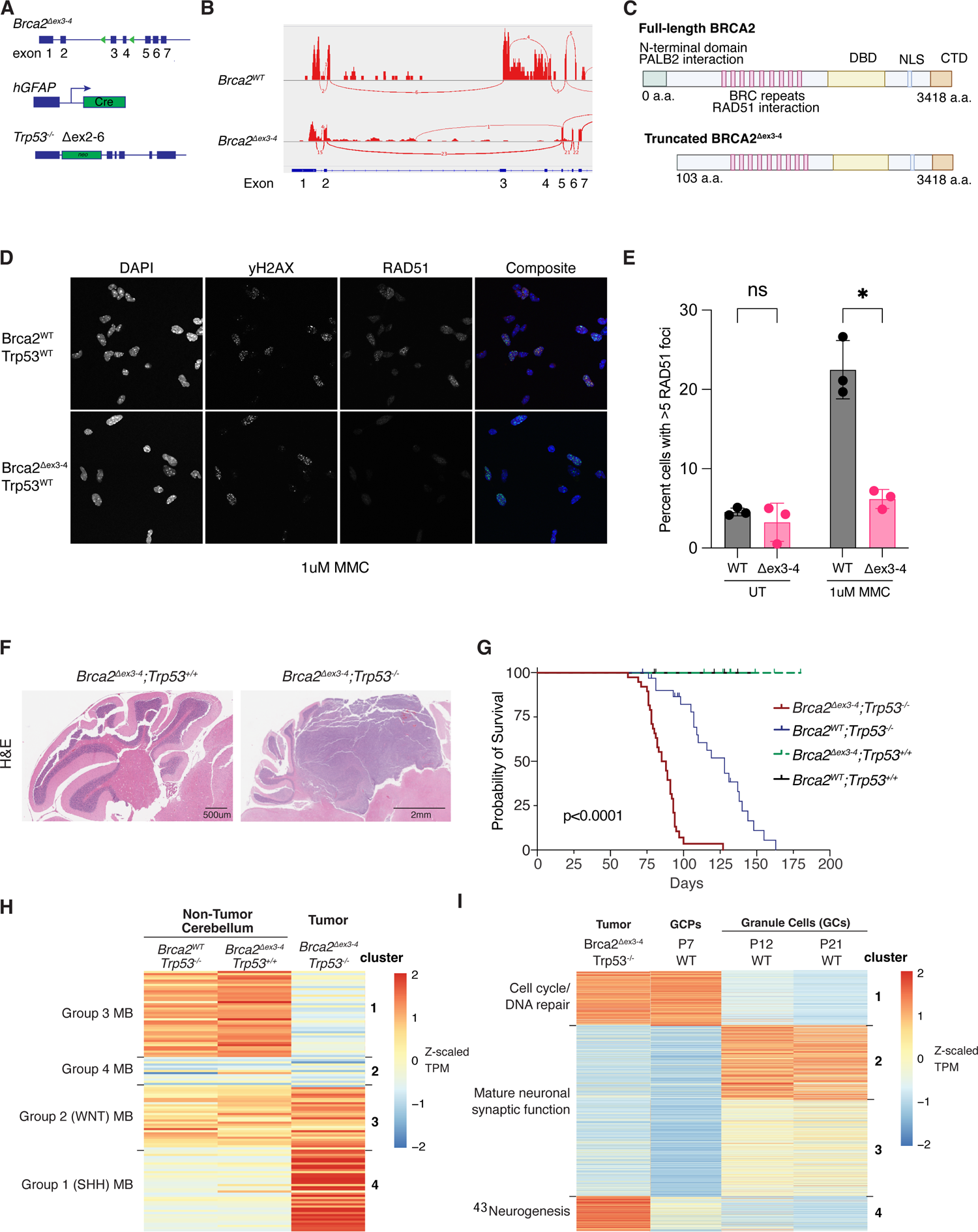
Loss of the PALB2-binding domain of *Brca2* and global loss of *Trp53* leads to the development of Sonic hedgehog subgroup medulloblastoma. (A) Summary of genetic perturbations in the mouse model of medulloblastoma usied in the study. *Brca2* with floxed exon 3-4, hGFAP-Cre transgene (FVB-Tg(GFAP-cre)25Mes/J), *Trp53* with neo cassette replacing exons 2-6 (B6.129S2-Trp53^tm1Tyj/J^). **(B)** Sashimi plots of splicing in *Brca2*^WT^ GCPs and *Brca2*^Δex3-4^ tumor demonstrates alternative splicing in *Brca2*^Δex3-4^ between exons 2 and 5. **(C)** Diagram of truncated BRCA2 without PALB2-binding domain. (D) *Brca2*^Δex3-4;^ *Trp53*^+/+^ GCPs are deficient in RAD51 foci formation after treatment with 1uM MMC for 5 hours. Representative images shown with DAPI (blue), yH2AX (green), and RAD51 (red). *Brca2^WT^; Trp53^+/+^* GCPs are competent for RAD51 foci formation. **(E)** Percentage of cells with greater than 5 RAD51 foci demonstrates an increase after MMC treatment for *Brca2*^WT^ GCPs but not *Brca2*^Δex3-4^ GCPs. (p=0.02). **(F)** Hematoxylin and eosin stained sagittal sections of mouse cerebellum from *Brca2^WT^; Trp53^-/-^* and *Brca2^Δex3-4^; Trp53^-/-^* mice at postnatal day 90 (P90) demonstrate medulloblastoma formation in the *Brca2^Δex3-4^; Trp53^-/-^* cerebellum. **(G)** Kaplan-Meier survival curve of *Brca2* and *Trp53* mutant animals demonstrates a significant decrease in survival due to tumor formation in *Brca2*^Δ^*^ex3-4^*; *Trp53*^-/-^ animals. **(H)** Heatmap of normalized gene expression data of medulloblastoma subgroup hallmark genes from RNA sequencing of bulk tumor and age-matched adult normal cerebellum. **(I)** Heatmap of normalized gene expression data from bulk tumor RNAseq as well as differentially expressed genes in proliferating GCPs compared to postmitotic granule cells (Matlik et al. 2023).

When *Brca2*^Δ^*^ex3-4^* was coupled with global *Trp53* knockout (Jacks et al. 1994), mice formed medulloblastomas with complete penetrance and exhibited a significant decrease in survival due to the tumor burden of double mutants (**Figure 1F**, G) in contrast to *Brca2^WT^; Trp53^-/-^* and *Brca2^Δex3-4^; Trp53^+/+^* animals, which did not develop MBs. This finding is consistent with MBs only forming on a *Trp53^-/-^* background in the majority of genetically engineered mouse models of medulloblastoma (Roussel and Stripay 2020). Although they did not form MBs, *Brca2^WT^; Trp53^-/-^* animals experienced increased mortality due to malignancies associated with Trp53 loss, such as lymphomas. MBs that formed in *Brca2^Δex3-4^; Trp53^-/-^*animals were highly proliferative and contained markers of DNA damage as indicated by the increased Ki67 and yH2AX staining, respectively (Supplemental Figure 1B). *Brca2^Δex3-4^; Trp53^+/+^* animals exhibited mild cerebellar hypoplasia at postnatal day 7—the developmental time at which GCPs are highly proliferative in the EGL (Supplemental Figure 1C). Tumors were easily identifiable by their EdU staining even at postnatal day 50 (Supplemental Figure 1D).

Guided by the previously identified expression changes in clinical subgroups of MB (Northcott et al. 2012; Kool et al. 2014; Miele et al. 2015) (Supplemental Table 1), we performed hierarchical clustering of tumor and age-matched control cerebellar expression data (**Figure 1H**). *Brca2*^Δ^*^ex3-4^*; *Trp53*^-/-^ MBs highly expressed genes associated with the SHH subgroup, including those involved in activated Hedgehog signaling and the cell cycle (Cluster 4, Supplemental Table 1) and downregulated genes associated with Group 3 and 4 MB including those associated with G-protein coupled receptor signaling, cell migration, and axon guidance (Clusters 1 and 2, Supplemental Table 1). We conclude that *Brca2*^Δ^*^ex3-4^*; *Trp53*^-/-^ MBs have a gene expression profile most like clinical SHH MB cases.

Previous studies have pointed to the cerebellar GCP as the cell of origin for SHH MB (Salsano et al. 2004; Hovestadt et al. 2019; Vladoiu et al. 2019). To determine how similar the tumor gene expression program was to GCPs, we performed hierarchical clustering using a list of genes found to be differentially expressed (padj < 0.01) in proliferating Math1-positive P7 GCPs compared to postmitotic NeuroD1-positive GCs (Matlik et al. 2023) (Supplemental Table 2).

Cluster 1 represents genes highly expressed in both proliferating, P7 GCPs, and tumors and associated with pathways of mitotic cell cycle, DNA replication, and homologous recombination. These include genes like *Cdk1* that regulates mitotic entry and *CyclinD2* that regulates G1 progression (**Figure 1I**). Other genes within cluster 1 are *Math1/Atoh1*, *Myc*, and *Pif1*. Clusters 2 and 3 were enriched in GCs at P12 and P21 and not expressed in tumors, and these clusters both contained genes associated with neurotransmitter signaling. Cluster 4 was the most tumor-enriched gene expression cluster, which diverged from non-tumor proliferating GCPs. This cluster included genes related to glial-guided migration such as *Sun1, Sun2,* and *Syne2* (Zhang et al. 2009), genes with roles in neurite outgrowth such as *Boc* and *Sema7a* (Vuong et al. 2017), and genes implicated in the cellular response to fibroblast growth factors (FGF), such as *Ctnnb1*, *Fgf3*, *Chrd*, and *Fgf5*. Overall, the gene expression analysis reveals that *Brca2*-deficient MBs have a gene expression program similar to P7 GCPs, but also acquire additional expression changes that may promote tumor growth through pathways divergent from those active during normal GCP proliferation.

### *Brca2*-mutant medulloblastoma is driven by *Ptch1* loss and characterized by mutations overlapping G-quadruplexes

To assess the mutational landscape of *Brca2^Δex3-4^* MB, we performed whole-genome sequencing on DNA isolated from four mouse medulloblastomas (MMB1-4) at 60x genome coverage and matched non-tumor forebrain tissue at 30x genome coverage. After filtering germline variants using the sequenced matched forebrain, we identified 643 variants across four tumors. The largest proportion of variants in the *Brca2^Δex3-4^; Trp53^-/-^* mouse MBs were small deletions **(Figure 2A, Supplemental Table 3, Supplemental Figure 2A)**. In addition, we detected structural variants (SVs) such as large deletions, translocations, inversions, and tandem-duplications that were highly variable in size, between 379 bp and 98 Mbp **(Supplemental Figure 2B).** A parallel analysis of copy number variation (CNV) identified 107 regions of copy number gain and 112 regions of copy number loss **(Supplemental Table 3, Supplemental Figure 2C)**.

**Figure 2.**
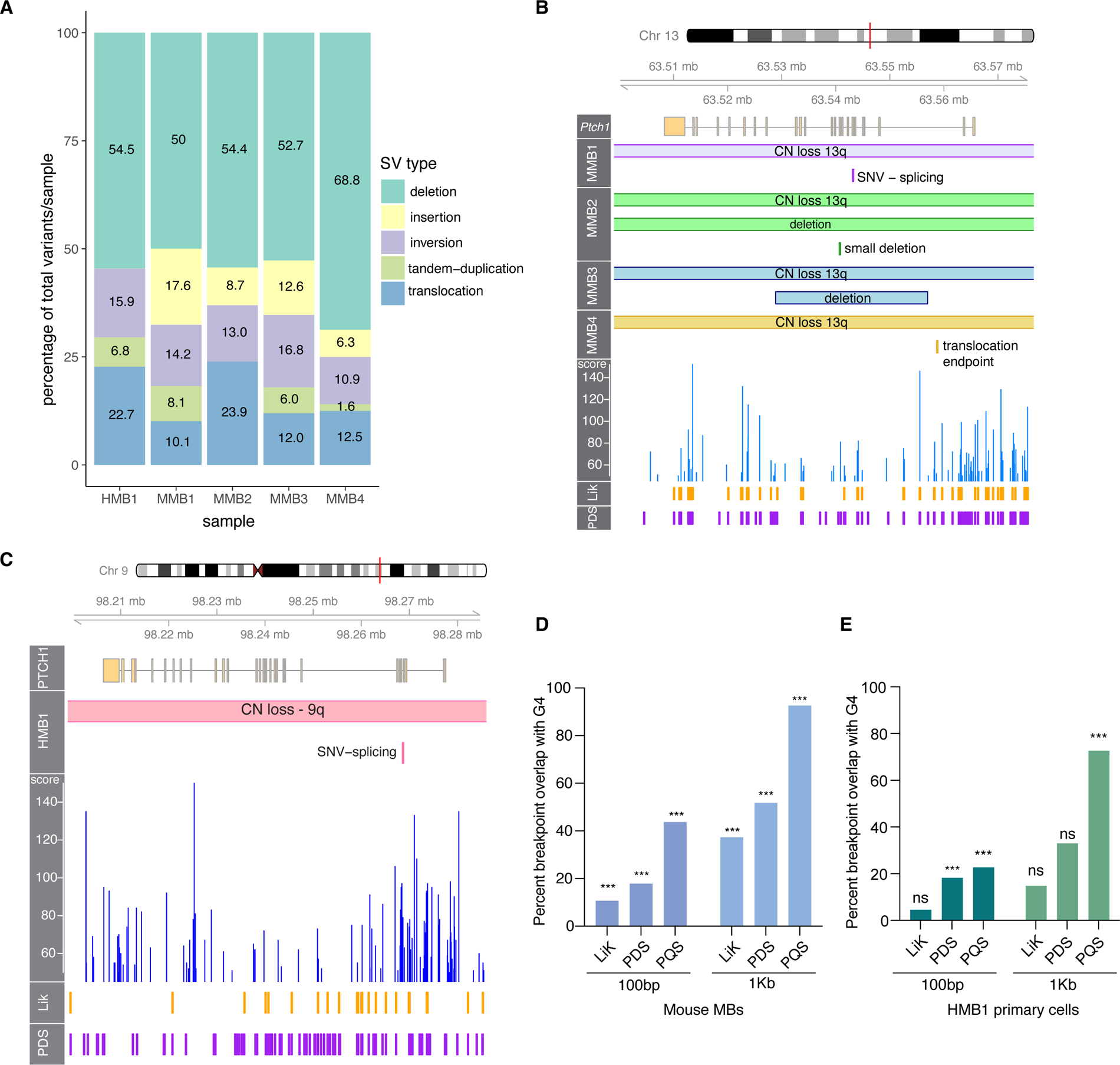
*Brca2* mutant medulloblastoma is driven by mutations that overlap with G-quadruplexes. **(A)** Percentages of structural variant class per sample of human MB primary cell line 1 (HMB1) and mouse MB samples 1-4 (MMB1-4). **(B)** Genome visualization (Gviz) plot of mouse *Ptch1* depicts CN loss of chromosome 13q and secondary mutation in each mouse MB sample in this G4-dense gene. **(C)** Gviz plot of HMB1 demonstrates CN loss of chromosome 9q and splicing SNV that inactivates *PTCH1*. **(D)** Statistical overlap test performed in regioneR (Gel et al. 2016) of tumor CNV breakpoints and G4-forming sequences from experimental and computational datasets (Hon et al. 2017; Marsico et al. 2019) demonstrates a significant overlap (p=0.001) of breakpoints and G4s when compared to random shuffling of breakpoint windows on the mouse genome. **(E)** Percentage overlap with PQSfinder G4s of each mutation type at 100bp breakpoints and 1kb breakpoints.

We also generated a primary MB cell line (HMB1) from an FA-D1 pediatric patient with germline compound heterozygous variants in *BRCA2* (c.9097_9098insT and c.9302 T>G) who was diagnosed with MB at the age of three years old. Clinically, this FA-D1 MB was determined to have SHH activation through increased YAP1 and GAB1 staining as well as TP53 deficiency by IHC. *TP53* variants detected through sequencing of HMB1 included a copy number loss of one allele and a pathogenic SNV (c.412G>C) in the second allele. The HMB1 primary cells expressed GCP markers including PAX6 and TAG1 (**Supplemental Figure 2D**). We sequenced the DNA isolated from this human primary MB cell line at 60x human genome coverage.

The HMB1 sample contained 44 structural variants, 27 regions of copy number gain, and 47 regions of copy number loss (**Supplemental Figure 2E-G, Supplemental Table 4**). These variants were determined after filtering using a panel of human controls due to the lack of normal tissue from this FA-D1 patient to perform germline filtering. Due to the inability to appropriately filter germline indel variants in HMB1, we omitted small indels from this analysis. SVs from the HMB1 line were enriched for deletions with a similar proportion as in the MMB samples (**Figure 2A**).

Few recurrent mutations were identified across the murine tumor samples, with the exception of mutations in *Ptch1* that were detected in all samples **(Figure 2B)**. Copy number loss of chromosome 13q where *Ptch1* is located was coupled in each sample with an inactivating mutation of the other allele of *Ptch1*. Mutations of the second allele of *Ptch1* included a single nucleotide variant predicted to lead to abnormal splicing and an early stop codon in MMB1, a large deletion and a smaller deletion in MMB2, a large deletion in MMB3, and a translocation upstream of *Ptch1* to chromosome 5 in MMB4. The human *BRCA2*-deficient MB carried *PTCH1* mutations following the pattern detected in mouse *Brca2^Δex3-4^* tumors. It contained a loss of one copy of chromosome 9q, a common copy number alteration in pediatric SHH MB, coupled with an inactivating SNV leading to alternative splicing and truncation of *PTCH1* **(Figure 2C**).

The *Ptch1* gene has previously been reported to have a high abundance of G-quadruplex forming sequences (Hoque et al. 2022). We utilized the putative quadruplex sequence (PQS) detection package pqsfinder (Hon et al. 2017) to identify the potential G4 sequences in mouse *Ptch1* (**Figure 2B**). pqsfinder computes a G4 score that indicates the predicted stability of each G4 and therefore its propensity to form. We found that *Ptch1* has in total 443 G4s, 88 of which are above score 50, indicating high predicted stability. Many of the somatic mutations in *Ptch1* occur between two peaks of high G4 stability coupled with high G4 density of overlapping G4s **(Figure 2B, 2C).**

Following the observation of a high G4 burden in the *Ptch1* locus, we hypothesized that other tumor variant breakpoints would significantly overlap with G4 loci in MBs. To test this hypothesis, we performed overlap testing of 100bp or 1Kb ranges around the breakpoint ends of the detected SVs and indels in the four mouse tumors against pqsfinder-derived putative G4s (**Supplemental Figure 3A-D**), as well as in two experimental datasets generated from stabilizing DNA libraries from mouse fibroblasts with lithium and potassium (LiK) or the G4-trapping drug pyridostatin (PDS) and performing mismatch analysis to identify where the sequencing polymerase stalled at stabilized G4s (Marsico et al. 2019). For both the 100bp and 1Kb breakpoint windows, we found high overlap of breakpoints with the PDS-stabilized or computationally predicted G4s and a more moderate overlap with the LiK dataset **(Figure 2D).** Permutation testing with regioneR (Gel et al. 2016) revealed that the overlaps were significant (p=0.001) compared to random shuffling of breakpoints in the genome **(Supplemental Figure 4A-F)** at both the 100bp and 1Kb breakpoint window sizes. We did not observe a particular bias for any class of mutation with G4 overlap **(Supplemental Figure 3A, B).** This indicates that the mutagenic repair of DNA breaks at G-quadruplexes may lead to diverse mutation outcomes.

We next performed overlap analysis using SV breakpoints from HMB1 utilizing pqsfinder-derived G4s identified from the human genome, and LiK and PDS-stabilized datasets detected from human HEK-293T cells (Marsico et al. 2019). Although there were only 88 SV breakpoints found in HMB1, we identified significant overlap with PQS G4s at both 100bp and 1Kb breakpoint windows from regioneR statistical tests **(Figure 2E, Supplemental Figure 4G-L).** As in mouse samples, we did not observe a bias for any SV type in the HMB1 sample (**Supplemental Figure 3E-G**. These data indicate that the presence of G4s may influence DNA breakage in human SHH-MB samples and lead to subsequent mutagenesis in G4-dense loci. The absence of BRCA2 may contribute to DSB formation at G4s or prevent proper repair leading to structural variant formation (see discussion).

### GCPs are highly replicative under SHH pathway activation and sensitive to G4 trapping by pyridostatin

The developmental biology of mouse GCPs can be assessed *ex vivo* by culturing GCPs as proliferative re-aggregates in suspension or as differentiated granule neurons in monolayer cultures plated on poly-D-lysine/Matrigel coverslips. GCPs have been shown to exhibit replication stress upon SHH pathway activation by treatment with the N-terminal fragment of SHH (Tamayo-Orrego et al. 2020). We confirmed this observation by treating wild-type GCPs with the Smoothened agonist (SAG) that leads to constitutive SHH pathway activation. The overall number of EdU-positive GCPs grown as re-aggregates increased with SAG treatment **(Figure 3A)**. To observe the influence of SAG on individual replication forks, we performed DNA combing on wild-type GCPs with sequential pulses of halogenated nucleotide analogs IdU and CldU **(Figure 3B)**. SAG treatment increased the average fork speed of wild-type GCPs from 1.2 kb/min in control cells to 2.3 kb/min, the highest of published fork velocity as measured by DNA combing across different cell lines (Techer et al. 2013), which range from 0.7 kb/min to 2kb/min.

**Figure 3:**
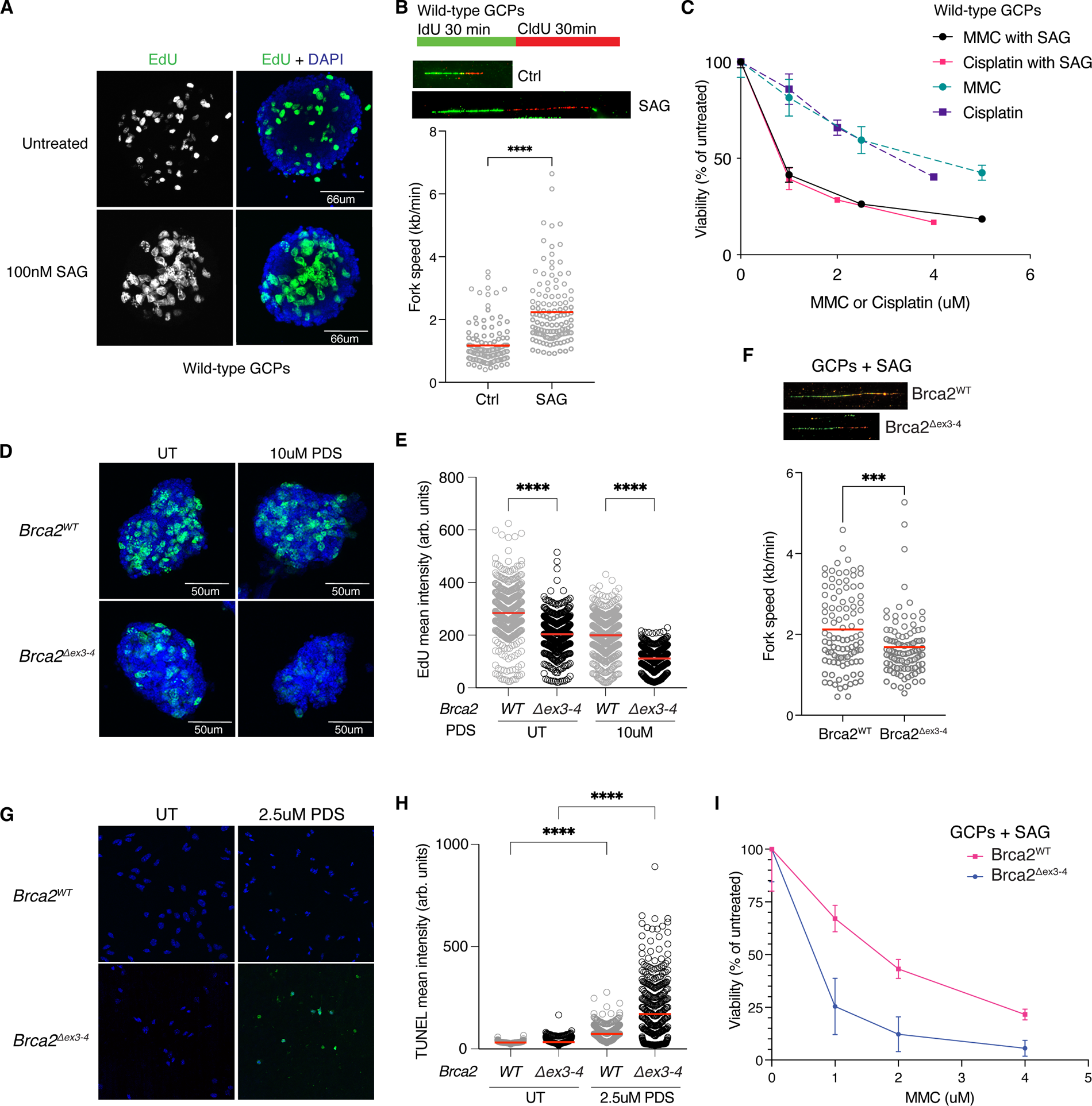
GCPs are highly replicative under SHH pathway activation and sensitive to G4 trapping by pyridostatin. **(A)** GCPs isolated from P7 wild-type C57BL.6 animals were cultured with or without 100nM SAG for 24 hours as proliferating re-aggregates and pulsed with EdU for 30 minutes. **(B)** SAG treatment for 24 hours additionally leads to faster replication fork speeds of 2.3kb/min on average, compared to 1.17 kb/min in non-SAG treated samples. **(C)** CellTiter-Glo viability assay on wild-type GCPs treated with Mitomycin C (MMC) and Cisplatin with or without SAG demonstrates a decrease in viability with SAG and treatment with a DNA crosslink-inducing agents. Error bars are +/- SD. **(D)** Representative confocal images of GCP re-aggregates treated with 100nM SAG and 10uM PDS for 2 hours prior to 10min EdU pulse. DAPI nuclear staining in blue and EdU staining in green. **(E)** Mean nuclear intensity of EdU incorporation in re-aggregates demonstrates a decrease in DNA replication level in *Brca2*^Δ^*^ex3-4^*; *Trp53^+/+^* GCPs after 2h of 10uM PDS treatment. 300 cells scored for each sample per replicate of three biological replicates (p<0.0001). **(F)** DNA combing of *Brca2*^Δ^*^ex3-4^*; *Trp53^+/+^*and *Brca2^WT^*; *Trp53^+/+^* GCPs in the presence of SAG. *Brca2*^Δ^*^ex3-4^* GCPs demonstrate a mildly decreased fork speed of 1.7kb/min compared to *Brca2^WT^*GCPs at 2.2kb/min (p=0.0005). **(G)** TUNEL staining for apoptosis in GCPs plated down immediately after isolation on PDL/Matrigel-coated coverslips demonstrates an increase in apoptosis in *Brca2*^Δ^*^ex3-4^*; *Trp53^+/+^* GCPs. DAPI nuclear staining in blue and TUNEL staining in green. 300 cells scored for each sample per replicate of three biological replicates (p<0.0001). **(H)** Mean intensity of TUNEL signal in individual adherent GCPs quantified in Imaris. Quantification demonstrates an increase in TUNEL-positive *Brca2*^Δ^*^ex3-4^*; *Trp53^+/+^* GCPs after PDS treatment (P<0.0001). **(I)** Cell viability measured with the CellTiter-Glo reagent demonstrates that survival of *Brca2*^Δ^*^ex3-4^*; trp53^+/+^ GCPs decreases more in the presence of MMC compared to *Brca2^WT^; Trp53^+/+^* GCPs. Error bars represent +/- SD.

We analyzed whether increased fork speed following SAG treatment led to greater DNA damage by examining yH2AX staining in proliferating GCPs. yH2AX intensity was significantly increased in SAG-treated GCPs, and this DNA damage was exacerbated further when cells were treated with the DNA crosslinking drug cisplatin which stalls replication **(Supplemental Figure 5A, B).** The observed increase in DNA damage also correlated with lower viability when wild-type GCPs were treated with SAG and crosslinking agents MMC or cisplatin, compared to DNA crosslinking drugs alone (**Figure 3C**). These findings indicate that SHH pathway activation by SAG drives increased replication speeds that render cells sensitive to replication stress. *In vivo*, GCPs require SHH secreted from neighboring Purkinje neurons (Wechsler-Reya and Scott 1999) to proliferate and expand to numbers of ∼60 billion in the human brain (Wagner et al. 2017). This suggests that during the massive proliferation that underlies neurogenesis, GCPs may exhibit an intrinsic vulnerability to replication-blocking DNA lesions that could be exacerbated upon SHH stimulation.

To test whether BRCA2 is required to promote genome stability in GCPs experiencing replication stress during the proliferative phase of cerebellar development, we performed DNA combing in *Brca2^Δex3-4^* GCPs. We found that fork speed decreased significantly in *Brca2^Δex3-4^; Trp53^+/+^* GCPs treated with SAG from 2.2kb/min to 1.7kb/min (**Figure 3F**). This slower fork speed could be due to increased stalling at genomic lesions in fast replicating cells when BRCA2 is absent. Based on our findings in tumors, G-quadruplexes could be the source of replication stress, thus we assessed whether G4s block replication in *Brca2^Δex3-4^* GCPs by treating proliferating GCP re-aggregates with pyridostatin (PDS), a G4 stabilizing chemical **(Figure 3D)**. Incorporation of 5-ethynyl-2′-deoxyuridine (EdU), a marker of replication, significantly decreased in *Brca2^Δex3-4^* GCPs after a two-hour treatment with high-dose PDS **(Figure 3E)**. The magnitude of the decrease was much smaller for *Brca2^WT^*GCPs indicating that BRCA2 deficiency reduces the ability of GCPs to replicated through stabilized G4s.

Cell viability downstream of replication stalling, as assessed by TUNEL staining for apoptosis following PDS treatment was decreased in *Brca2^Δex3-4^; Trp53^+/+^*, but not in *Brca2^WT^; Trp53^+/+^* GCPs when treated with PDS **(Figure 3G, H).** These data indicate that BRCA2 deficiency sensitizes GCPs to cell death after treatment with G4-stabilizing drugs, perhaps due to an increase in DNA breaks at unprotected stalled forks or due to deficiency of DSB repair. As predicted, treatment with DNA crosslinking agents MMC and SAG also reduced *Brca2^Δex3-4^; Trp53^+/+^* GCP viability **(Figure 3I),** as expected due to BRCA2 function in interstrand crosslink repair.

### The PIF1 helicase protects *Brca2*^Δ^*^ex3-4^* medulloblastoma cells against genomic instability at G-quadruplexes

The potential importance of G-quadruplex resolution in the BRCA2 deficient medulloblastoma was further highlighted when we assessed gene expression changes in tumor compared to age-matched normal cerebellum and identified that the 5’-3’ DNA helicase PIF1, which has affinity for and promotes processivity through G4s (Paeschke et al. 2011; Byrd and Raney 2017) was one of the most upregulated genes in tumors **(Figure 4A)**. Topoisomerase IIA (*Top2A*), which functions in releasing DNA supercoiling during replication, was also highly upregulated in tumor samples, highlighting the particular importance of DNA processing during replication in MBs. Notably, other G4 helicases such as *Blm*, *Wrn*, and *Brip1/Fancj* were also expressed but not differentially expressed in tumor versus normal cerebellum. RNAseq from samples serially harvested across tumor development, by sorting EdU-pulsed cells from the adult, postmitotic cerebellum at P35, P50, and P75, revealed that *Pif1* upregulation occurred as early as P35, which coincided with early tumor development (**Figure 4A, Supplemental Figure 1D**).

**Figure 4:**
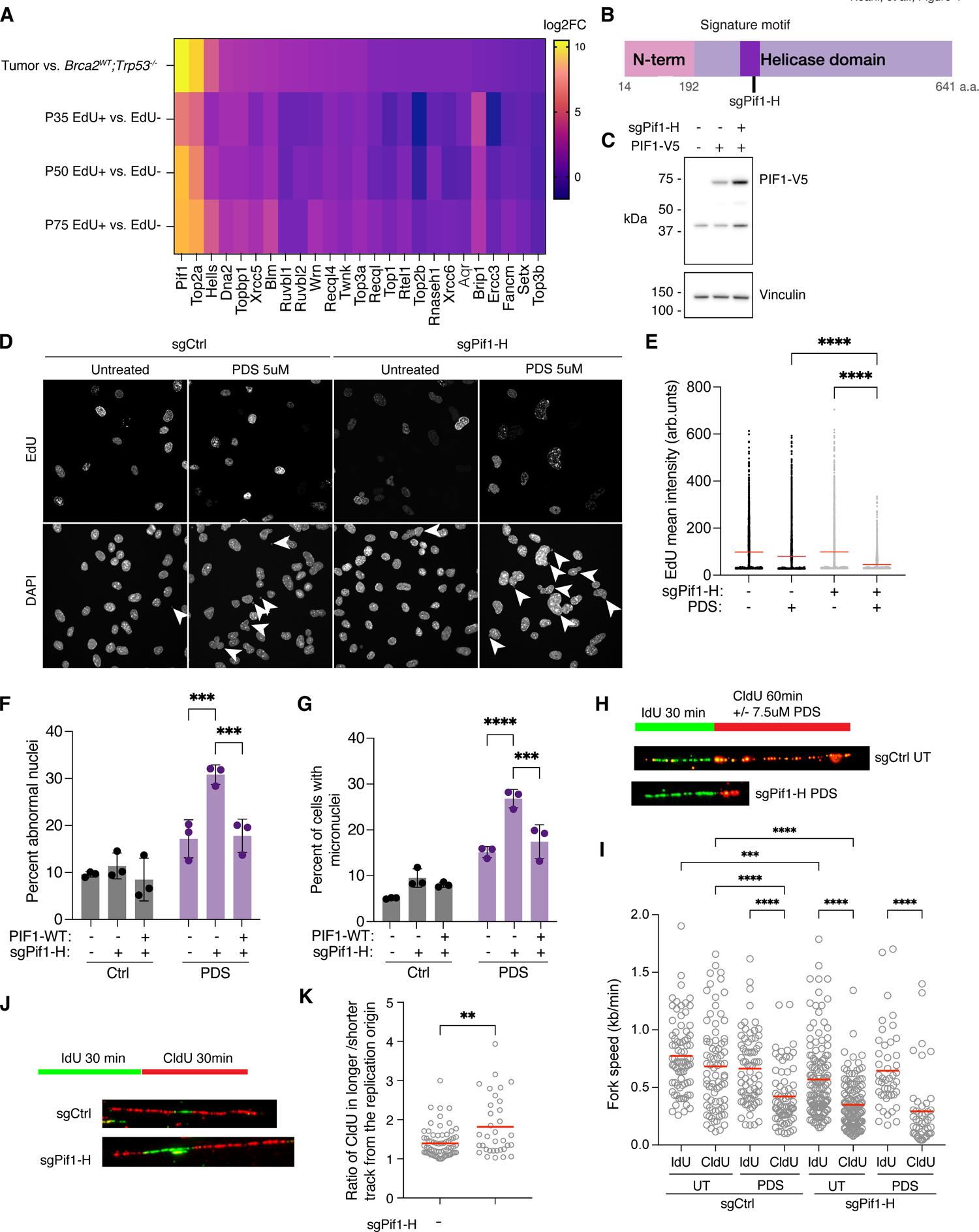
The PIF1 helicase protects *Brca2*^Δ^*^ex3-4^* medulloblastoma cells against genomic instability at G-quadruplexes. **(A)** Heatmap of log2 fold change (log2FC) values from differential gene expression datasets demonstrates *Pif1* and *Top2a* upregulation in *Brca2*^Δ^*^ex3-4^* tumor compared to non-tumor *Brca2*^WT^ cerebellum. **(B)** Schematic of PIF1 protein indicating the domain targeted by the sg*Pif1*-H in primary *Brca2*^Δ^*^ex3-4^* tumor cells. **(C)** Immunoblot with an antibody recognizing V5-tagged PIF1 and the loading control (Vinculin) demonstrates an increase in PIF1 expression when endogenous *Pif1* is knocked out. **(D)** Representative images of sgNeg or sgPif1-H KO cells stained with DAPI and EdU after 48h of treatment with and without 5uM PDS. White arrows point to abnormal nuclei and micronuclei, which increase in bulk sgPif1-H KO with PDS treatment. **(E)** Mean nuclear intensity of EdU demonstrates a decrease in the level of DNA replication in sgPif1-H MB cells (p<0.0001). **(F)** Abnormal nuclei percentage as determined with a Imaris machine-learning pipeline. 300 nuclei scored per sample in each biological replicate of three biological replicates (p=0.0006, p=0.0009). **(G)** Micronuclei hand-counted per 300 DAPI-stained cells in each sample across three biological replicates (p=0.0001, p=0.0009). sgNeg and sgPif1-H KO cells demonstrate increased micronuclei after PDS treatment, with a greater increase in sgPif1-H KO cells. **(H)** DNA combing scheme for MB cells where cells are pulsed with IdU for 30min, followed by a 60min CldU pulse with or without PDS to assess fork stalling after G4 stabilization. **(I)** Fork speed quantification reveals a decrease in fork speed in sgCtrl MB cells with PDS treatment, untreated sgPif1-H cells as replication progresses in the second CldU label, and sgPif1-H cells treated with PDS (p<0.0001). **(J)** Schematic of labeling to assess replication speed asymmetry. **(K)** Combing of MB1 cells demonstrating greater origin asymmetry when Pif1-H guide was used (p=0.004). UT=untreated.

Beyond its ability to unwind G-quadruplex DNA, DNA/RNA hybrids, and forked DNA (Chib et al. 2016; Li et al. 2016; Tran et al. 2017; Pohl and Zakian 2019), PIF1 has been shown to facilitate genome stability of G4-rich DNA in yeast (Paeschke et al. 2011; Paeschke et al. 2013) and RAS-transformed human fibroblasts (Gagou et al. 2011; Gagou et al. 2014). In *Drosophila* embryos, PIF1 was found to be necessary for overcoming replication stress and its loss was synthetically lethal with BRCA2 mutation (Kocak et al. 2019).

To assess the role of PIF1 in *Brca2^Δex3-4^* MB, we utilized primary cells isolated from mouse MBs. These cells had neuron-like morphology, remained proliferative in culture, and expressed markers typical of *Brca2^Δex3-4^* tumors such as *Tag1* and *Pax6* **(Supplemental Figure 6A)**. We targeted PIF1 with a CRISPR/Cas9 sgRNA directed at the signature motif within the helicase domain required for PIF1 activity (Geronimo et al. 2018) (sgPif1-H). Prior to CRISPR-directed cutting, primary MB cells were complemented with C-terminally V5-tagged mouse PIF1 cDNA with a synonymous PAM-site mutation making it resistant to CRISPR guide directed cleavage **(Figure 4B)**. The gRNA produced high bulk cutting efficiency within PIF1 **(Supplemental Table 5)** and the V5-tagged PIF1 expression was maintained after cutting using sgPif1-H **(Figure 4C).**

Single-cell clones with frameshift mutations in the PIF1 helicase domain, exhibited significant increases of large, abnormal nuclei and micronuclei **(Supplemental Figure 6B-D).** These phenotypes of genomic instability were exacerbated further when clones were complemented with a helicase dead mutant of PIF1 (E316Q) **(Supplemental Figure 6E-G)** (Zhou et al. 2016). Bulk targeting with sgPif1-H resulted in increased levels of aberrant nuclei that were further increased following PDS treatment but decreased with complementation using PAM-resistant WT PIF1 **(Figure 4D, 4F).** Similarly, micronuclei were found to be significantly more frequent in PIF1-deficient cells treated with PDS (**Figure 4G**). They also exhibited decreased EdU incorporation, indicating slowed replication rates consistent with replication stress **(Figure 4E).**

We then examined the effects of PDS on individual replication forks of PIF1-deficient MB cells by DNA combing. PIF1-deficient MB cells had slower rates of replication than control cells, with particularly decreased fork speed during the second pulse, which suggests that these forks encounter stalling lesions (**Figure 4H, I**). Further support for heightened fork stalling in these cells came from analysis of replication fork asymmetry, which was significantly increased in PIF1-KO cells (**Figure 4J, K**). PIF1-competent MB cells also demonstrated a decrease in fork speed after PDS treatment, which indicates that endogenously expressed PIF1 can be overwhelmed by high levels of stabilized G4s. These findings indicate that replication through G4 DNA is facilitated by the PIF1 helicase in primary MB cells and suggest that PIF1 upregulation is essential to support replication through G4s as *Brca2^Δex3-4^* MB cells proliferate. Targeting PIF1 by chemical inhibition or genetic modification in BRCA2-deficient MBs could allow for selective killing of tumor cells, sparing surrounding normal cerebellum that lacks *Pif1* expression.

## Discussion

This study aimed to identify endogenous causes of vulnerability of a highly proliferative developmental cell population, the cerebellar granule cell progenitor, in the setting of BRCA2-deficiency. Using *in vitro* and *in vivo* approaches, we have identified G-rich DNA capable of creating G4s, as being a challenge for BRCA2-deficient GCPs. We determined a significant association of structural variant and indel breakpoints with G4s in the spontaneously formed medulloblastomas of the *Brca2^Δex3-4^; Trp53^-/-^* mouse model. These included inactivating mutations in the G4-dense gene *Ptch1,* a most likely primary driver of tumorigenesis in all MBs formed in our model, which is consistent with previous studies of a BRCA2 model of MB (Frappart et al. 2007; Frappart et al. 2009), genomic studies of SHH-group MB from a large patient cohort (Kool et al. 2014) and rare FA-D1 patients (Waszak et al. 2018) that also identified *PTCH1* mutation as recurrent in pediatric SHH tumors.

Prior studies suggest that different tissues and cells may display vulnerability to different endogenous sources of genome instability (Langevin et al. 2011; Garaycoechea et al. 2012; Dingler et al. 2020; Jung et al. 2022). It is unclear why the GCPs would be particularly vulnerable to instability at G4s. Both replication speed and DNA damage increase when GCPs are stimulated by SHH (Tamayo-Orrego et al. 2020). GCPs therefore may be particularly vulnerable to fork stalling because of the demands of their postnatal expansion. Under those conditions, BRCA2 plays an integral role in preventing genomic instability at G4s in GCPs, averting medulloblastoma development. However, it remains to be determined if the essential function of BRCA2 at G4-rich regions is to protect stalled forks allowing for its non-mutagenic processing and prevention of replication fork collapse, or alternatively, the homology-directed repair of the DNA damage that occurs when such collapse occurs at the G4s.

Given that that embryonal tumors in patients with biallelic pathogenic variants of BRCA2 include neuroblastoma and Wilms tumors, it is tempting to speculate that the essential function of BRCA2 in prevention of these two blue cell tumors will also include BRCA2 function at G4 quadruplexes. This hypothesis will have to be tested in future studies. The connection of G4s and BRCA2 may also be important in other settings of BRCA1/2 deficiency, as the stabilization by PDS leads to cell death in BRCA1/2 deficient cells (Zimmer et al. 2016) and in patient derived xenografts (Groelly et al. 2022). More broadly, CNV breakpoints in human cancers were previously found to be located in proximity to G4s (De and Michor 2011), with G4-forming sequences located downstream of CNV loci potentially causing leading strand replication stalling and DNA breakage leading to mutagenesis.

G-quadruplexes may not be the only lesions that are being tended by BRCA2 in the context of GCPs. Reactive aldehydes, including formaldehyde and acetaldehyde, are formed during cellular metabolism and are detoxified by the enzymes ALDH2 and ADH5 (Dingler et al. 2020). High levels of reactive aldehydes overwhelm the detoxification capacity of ALDH2/ADH5 and result in DNA interstrand crosslinks (ICLs) (Garaycoechea et al. 2018; Hodskinson et al. 2020) that require functional Fanconi anemia pathway for repair (Langevin et al. 2011). Given the high metabolic demands of postnatal GCP development, the impact of aldehyde accumulation on tumorigenesis in *Brca2*-deficient GCPs should be considered.

BRCA2 has also been shown to regulate the accumulation of RNA/DNA hybrids or R-loops. It potentially does so by regulating promoter-proximal pausing through interaction with RNA polymerase II (Shivji et al. 2018), through interaction with the TREX-2 mRNA export complex (Bhatia et al. 2014), and during replication by interacting with the transcription regulator ZFP281 (Wang et al. 2022). R-loops were also observed to increase in breast cancer cells following estrogen treatment which may contribute to DNA replication stress and genomic rearrangements at R-loop-forming loci (Stork et al. 2016). Future studies should examine the R-loop burden and localization specifically in GCPs, to determine if they may contribute to their genomic instability if BRCA2 is absent. As the G-quadruplexes are often colocalized with R-loops, especially at G-rich regions that experience negative supercoiling and are actively transcribed (Duquette et al. 2004), future G4 mapping in combination with R-loop mapping in GCPs could highlight more specific regions of vulnerability that would also consider the transcriptional landscape of GCPs.

Analysis of the transcriptome of medulloblastomas that developed in *Brca2^Δex3-4^; Trp53^-/-^* mouse model hinted at the importance of *Pif1* in tumor cells. We found that PIF1 was expressed both in tumor cells and proliferating GCPs, but not expressed in the normal adult cerebellum. This finding nominates PIF1 as a potential therapeutic target in BRCA2-deficient MB. PIF1 loss led to decreased replication speed in otherwise unperturbed primary tumor cells, indicating that loss or inhibition of this helicase may stall replication forks at lesions such as G4s and could reduce proliferation of tumor cells.

PIF1 overexpression has been associated with other cancers. Testicular, ovarian, and breast cancers express PIF1 at high levels (Park et al. 2019). High PIF1 expression was found to be correlated with reduced survival outcomes in neuroblastoma patients (Chen et al. 2020), indicating that the role of PIF1 in G4 biology may be relevant in other cancers.

Studies in model organisms have also shown the importance of G4 resolution in maintaining genome stability. In *C. elegans*, loss of the *dog1/FANCJ* G4-unwinding helicase led to both small deletions and more complex rearrangements upstream of G4 loci (Cheung et al. 2002; Kruisselbrink et al. 2008). FANCJ was shown to directly resolve G4s and prevent replication stalling that was enhanced when G4s were stabilized (Castillo Bosch et al. 2014; Sato et al. 2021). It remains to be tested if PIF1 and FANCJ act redundantly to prevent instability at G4s in GCPs and MB cells.

To conclude, our study suggests that BRCA2 and G4 helicases work in concert in GCPs to prevent the accumulation of mutations at G4 loci in this highly proliferative neuronal progenitor population that has an intrinsic vulnerability to DNA damage during replication. BRCA2-deficient MB cells were also shown to upregulate G4 helicases to foster tumor cell proliferation.

## Methods

### Mouse husbandry and tumor monitoring

All mouse experiments were approved by the Rockefeller University Institutional Animal Care and Use Committee (IACUC protocol #23039-H). *Brca2^Δex3-4^* mice (a gift from Dr. Maria Jasin) were crossed with *Trp53^+/-^*B6/129S mice (B6.129S2-Trp53tm1Tyj/J, Jackson Laboratories) and *hGFAP-Cre* mice (FVB-Tg(GFAP-cre)25Mes/J, Jackson Laboratories) to generate *Brca2^Δex3-4^* in the CNS coupled with global *Trp53* knockout. Double-knockout animals that form medulloblastomas were checked bi-weekly for cerebellar tumor formation with the IVIS ultrasound system in the Rockefeller University Comparative Bioscience Center. All mice were monitored for weight loss, doming of the head, ataxia, or failure to thrive, and sacrificed humanely. 100 animals were monitored for the survival and tumor outcome cohort. At the humane euthanasia endpoints, tumors were collected and dissociated into primary tumor cell lines, flash frozen for RNA and DNA isolation, or whole brain dissection and fixation for histology. Mice were genotyped using primers in **Supplemental Table 6**, using GoTaq Green Master Mix (Promega #M7122) for *Brca2, Trp53,* and *hGFAP-Cre* alleles.

### Isolation of primary medulloblastoma cells

Primary medulloblastoma tumor cells were isolated as previously described in (Badodi et al. 2019). Briefly, after mice were humanely sacrificed, friable tumors were dissected from the cerebellum and dissociated with Accutase for 5min at 37C. Tumors were dissociated in Accutase with pipetting and were spun down for 5min at 500g. Tumor cells were plated on tissue-culture 6-well plates pre-coated in 10ug/mL laminin (Sigma L2020) in NeuroCult Mouse/Rat Proliferation Kit medium (STEMCELL Technologies #05702) with 1x Pen/Strep, epidermal growth factor (EGF, Peprotech #AF-100-15), basic fibroblast growth factor, recombinant human (bFGF, Peprotech #100-18B), and heparin (STEMCELL Technologies, #07980).

### FA-D1 Patient Medulloblastoma Primary Cells

FA-D1 patient F89P1 was entered into the Rockefeller University International Fanconi Anemia Registry (IFAR) under IRB number AAU-0112. Medulloblastoma tissue was collected after surgical resection and primary cells were isolated in the same manner as for mouse MBs as described below. Primary human MB cells (HMB1) were cultured in NeuroCult NS-A Proliferation Kit (Human, STEMCELL Technologies #NC0668437) with 1x Pen/Strep, epidermal growth factor (EGF, Peprotech #AF-100-15), basic fibroblast growth factor, recombinant human (bFGF, Peprotech #100-18B), and heparin (STEMCELL Technologies, #07980).

### Isolation of primary GCPs

GCPs were isolated at P7 from C57BL/6 (strain #000664, Jackson Laboratories) wild-type or *hGFAP-Cre; Brca2^Δex3-4^; Trp53* mutant animals as previously described (Hatten 1985). Cerebella were dissected in ice-cold CMF-PBS and dissociated with trypsin-DNAse I for 5min at 37C. Dissociated cerebella were centrifuged for 5min at 700g at 4C. Trituration of cerebellar tissue was performed in CMF-PBS with DNase I with fine fire-polished Pasteur pipets followed by extra-fine fire-polished Pasteur pipets. GCPs were enriched using the Percoll separation gradient. The GCP-containing small cell fraction was collected and pre-plated on a Petri dish for 30min followed by one hour on a tissue culture-treated dish, allowing for plating of fibroblasts and other contaminating cells. After preplating, GCPs were pelleted at 700g for 5min at 4C and counted prior to culturing.

### Culture of GCPs on coverslips

GCPs were cultured on glass coverslips pre-coated with poly-D lysine (0.1mg/mL, Sigma #P1024) for 1h followed by drying and Matrigel coating (growth factor-reduced, Corning #354230). GCPs were cultured in Basal Medium Eagle (Gibco #21010-046) with 2mM L-glutamine (Gibco #25030-016), 1X Pen-Strep (Gibco #15140-015), 0.9% glucose (Sigma #G8769), 10% horse serum (Gibco #16050-122, heat-inactivated) and 5% fetal bovine serum (Gibco #26140-079, heat-inactivated)) at a final concentration of 50,000 cells per coverslip in 500ul media. GCPs were treated with 100nM SAG (Cayman Chemical, #11914) for 3h or 12h as well as pyridostatin (MedChemExpress #HY-15176A) at 2.5uM, 5uM, or 10uM. EdU (ThermoFisher #E10187) was added at 10uM 30min prior to the end of each timepoint.

### Culture of re-aggregate GCPs in suspension

GCPs were plated at 2*10^6^ cells in 500ul of media in Ultra Low Attachment 24-well plates (Corning #3473) or 100,000 cells in 100ul in opaque-bottom 96-well plates (ThermoFisher #15042) and allowed to re-aggregate for 16h prior to treatment with 100nM SAG, mitomycin C, cisplatin, or pyridostatin. Re-aggregate GCPs grown in 24-well plates were pulsed with 10uM EdU at the end of the treatment window and fixed in 4% methanol-free PFA for 15min prior to being centrifuged onto slides with the ThermoFisher Cytospin 4 Cytocentrifuge at the Rockefeller University Flow Cytometry Resource Center. Re-aggregate GCPs grown in 96-well plates were processed for cell viability readings in CellTiter-Glo as described below.

### Generation of Pif1-KO MB Cells

CRISPR RNP cutting was performed with Cas9 complexed with an assembled sgRNA using the IDT Pre-Designed Alt-R CRISPR/Cas9 system. mPif1 guides “AC” and “AA” were selected from the gRNA design tool due to high specificity and predicted cutting efficiency (Mm.Cas9.PIF1.1.AA -“sgPif1-N” and Mm.Cas9.PIF1.1.AC - “sgPif1-H”). gRNAs were complexed in oligo duplex buffer (IDT) with a tdTomato-labeled tracrRNA (IDT #1075927) prior to complexing with SpCas9 V3 (IDT #1081059). CRISPR-RNP complexes were delivered to primary medulloblastoma cells using the P3 Primary Cell Nucleofection Kit (Lonza #V4XP-3032) for the 4D Nucleofector X (Lonza #AAF-1003X). Program optimization was performed with pMaxGFP plasmid included in the nucleofection kit and the program CL-133 was found to be the optimized protocol for primary MB cells. CRISPR cutting efficiency was determined with Synthego ICE analysis using 700bp of sequenced PCR around the cut site for each guide.

### Western blotting

Protein lysates were generated by sonicating and boiling 1*10^6 cells in 100ul of 2X Laemmli Buffer (Bio-Rad) with BME. 20ul of lysate was added to the wells of precast 10-well 4-12% Bis-Tris PAGE gels in MOPS running buffer. After overnight transfer, membranes were blocked in 5% milk in TBST (10mM Tris-Hcl pH 7.5, 150mM NaCl, 0.1% Tween-20) and incubated in primary antibodies overnight at 4C with rocking (**Supplemental Table 7**). Membranes were washed in TBST 3x5min before being incubated with HRP-conjugated secondary antibodies for 1h at RT with rocking. Membranes were washed again in TBST and detected by enhanced chemiluminescence. Western blots were visualized with the Azure c300 imaging system.

### Immunofluorescence

Coverslips were fixed in 4% paraformaldehyde in PBS for 15min at RT. Permeabilization and block were performed in 3% BSA in PBS with 0.5% Triton-X for 30min at RT with rocking. Coverslips were then incubated with primary antibodies diluted in 3% BSA overnight at 4C (**Supplemental Table 7**). Washes were performed in PBS 3x5min. Secondary antibodies were diluted in 3% BSA in PBS and incubated for 1h at RT and washes were performed 3x5min in PBS. Hoechst 33342 (ThermoFisher) was used to stain DNA and coverslips were incubated in 1:2000 diluted Hoechst in PBS for 20min at RT at the end of the wash. Coverslips were mounted on SuperFrost Plus slides in Fluoromount-G and allowed to cure overnight. Imaging was performed on an LSM 880 confocal microscope at the Rockefeller University Bio-imaging Resource Center.

### EdU Staining

Cells pulsed with 10uM EdU for 30min were washed in 1x PBS and fixed in 3.7% formaldehyde in PBS for 15min at RT. Cells were stained with Click-iT™ EdU Alexa Fluor™ 488 Imaging Kit (Invitrogen, C10337) according to manufacturer’s protocol. DNA was stained with Hoechst 33342 and coverslips were mounted in Fluoromount-G.

### Image analysis with Imaris

Confocal Z-stacks were analyzed using Imaris imaging software. DAPI was used to mask nuclear surfaces and to separate touching surfaces based on the average diameter of nuclei. To quantify the number of abnormal nuclei in images, a machine learning pipeline was created using Imaris’s machine learning filter on DAPI-masked surfaces (**Supplemental File 1**). 200 normal and 200 abnormal nuclei were selected to train the machine learning pipeline and quantification of the trained pipeline was validated in a subset of training samples with hand-scoring. On subsequent datasets, the machine learning filter was run in an automated way to reduce bias in image quantification. Within each nuclear surface, mean intensity values for the other immunofluorescence channels such as yH2AX and EdU were also automatically quantified and exported.

### Image Analysis in FIJI

Micronuclei were hand-scored in FIJI by first counting the total number of nuclei (at least 300 nuclei per three biological replicates) with the CellCounter plugin and then counting cells with at least one micronucleus associated in the immediate vicinity of the nucleus. Micronuclei were reported as percent of cells containing one micronucleus. Due to the low cytoplasmic volume of MB primary tumor cells, micronuclei were clearly associated with the nucleus. RAD51 foci were additionally hand-scored, with total nuclei counted with the CellCounter plugin followed by counting of cells with greater than 5 bright RAD51-positive foci. 100 cells were scored per three biological replicates for this assay.

### DNA combing coverslip production

Silanized coverslips were produced as described in (Gallo et al. 2016; Moore et al. 2022) with the modification of plasma cleaning coverslips with the Gatan Model 950 Advanced Plasma System with atmospheric air for 10 minutes.

### DNA combing

Cells were labeled with IdU (100uM) and CldU (100uM) for the specified pulse times while in exponential growth. Cells were harvested in Accutase and washed in PBS prior to resuspension in 45ul combing Resuspension Buffer (PBS with 0.2% sodium azide). 45ul of 2% low melt agarose was added to briefly warmed cell suspension and poured into agarose plug molds. Agarose plugs were digested overnight at 55°C in 1mg/mL proteinase K, 1% N-Lauroylsarcosine, 0.2% sodium deoxycholate, 100mM EDTA, 10mM Tris-HCl, pH 7.5. Plugs were wash three times for at least an hour in 1X TE pH 8.5 with 100mM NaCl. Plug melting was performed in 1mL freshly prepared combing buffer (0.5M MES pH 5.5) for 68°C for 20 minutes followed by overnight incubation with 1ul beta-agarase at 42°C. Combing was performed onto silanized coverslips made in house using the Molecular Combing System (Genomic Vision). Slides were dried for 2h at 65°C and denatured in 0.5 M NaOH + 1M NaCl for 8min or moved directly into YOYO-1 DNA stain (ThermoFisher) for quality control check. Denatured combed coverslips were dehydrated in 70%, 90%, and 100% ethanol for 5min sequentially and dried in the dark at RT. Blocking was performed at RT for 1h in 5% FBS in PBS and primary antibodies were incubated overnight diluted in 5% FBS in PBS. Coverslips were washed 3x5min in 5% FBS in PBS prior to 1h secondary antibody incubation at RT. Coverslips were mounted in Fluoromount-G and imaged on an Inverted Olympus IX-71 DeltaVision (Applied Precision) microscope. Combed DNA fibers were scored in FIJI software.

### Cell viability assay using Cell Titer-Glo

100,000 GCPs per well were grown in opaque-bottom 96-well plates in triplicate and were processed by the addition of 1:1 CellTiter-Glo reagent directly to culture medium after 48-72 hours of treatment. Plates were agitated for 30min at RT to allow cell lysis to occur prior to reading luminescence values on the BioTek Synergy Neo2 microplate reader in the Rockefeller Drug Discovery Resource Center.

### RNA-sequencing

Total RNA was isolated using the Zymo Quick-RNA Miniprep Kit with on-column DNA digestion. mRNA was enriched with the NEBNext Poly(A) mRNA Magnetic Isolation Module (NEB #E7490) and library preparation was performed with the NEBNext Ultra II RNA Library Prep Kit for Illumina (NEB #E7770S). Indices used were the NEBNext Multiplex Oligos for Illumina (NEB #E7335, #E7500). Sequencing libraries were evaluated on the Agilent 2200 TapeStation with D1000 High Sensitivity ScreenTape and sequenced on a NextSeq 500 High Output sequencer to generate 75bp reads at the Rockefeller University Genomics Resource Center. Transcript abundance was determined using Salmon (v0.8.1) and the reference transcript sequences from TxDb.Mmusculus.UCSC.mm10.knownGene (v3.4.0) (Patro et al. 2017). Transcript counts from Salmon were imported into R with the tximport R Bioconductor package (v1.20), and differentially expressed genes were determined with the DESeq2 R Bioconductor package (v1.20) (Love et al. 2014). Significant genes were considered as p-adj < 0.01 and log2FC ≥ 1. GO gene set enrichment analysis (GSEA) was performed in clusterProfiler (v4.0.5) (Yu et al. 2012). Heatmaps were generated in pheatmap (v1.0.12) (Kolde 2019).

### Illumina whole-genome sequencing

DNA extraction was performed on 4 flash frozen mouse medulloblastomas and matched forebrain normal controls using UltraPure phenol:chloroform:isoamyl alcohol (25:24:1) (ThermoFisher 15593031). Frozen tumor tissue was homogenized with bead-beating using the Qiagen TissueLyser II in Qiagen DNeasy Blood and Tissue Kit Buffer ATL with proteinase K added. Illumina WGS library preparation and sequencing was performed at the National Institutes of Health (NIH) Intramural Sequencing Center using the Illumina PCR-free TruSeq library preparation kit. Tumor samples were sequenced to 60x coverage of the genome and matched normal samples were sequenced to 30x genome coverage.

### Whole-genome sequencing alignment and mutation calling

Sequencing data from mouse was aligned to NCBI mouse reference genome GRCm38 (mm10) and sequencing data from the HMB1 primary human MB line was performed using GRCh37 reference genome (hg19) using BWA-MEM2 (https://github.com/bwa-mem2/bwa-mem2). Duplicate reads were identified with (https://github.com/samtools/samtools). Somatic SNVs were called with CaVEMan (https://github.com/cancerit/CaVEMan) and somatic indels called by Pindel (https://github.com/cancerit/cgpPindel). SVs were called with BRASS (https://github.com/cancerit/BRASS) and further annotated by AnnotateBRASS (https://github.com/MathijsSanders/AnnotateBRASS) as described previously (Moore et al. 2020). After variant calling, filtering of artefacts, alignment errors, and low-quality variants was done as described previously (Moore et al. 2020; Webster et al. 2022) followed by filtering of sample-matched normal forebrain controls for mouse samples. For the human MB primary cell line HMB1, which lacked a paired normal control, variant filtering was performed with an in-house human panel of controls (Webster et al. 2022). CNV regions were determined with CNVkit (Talevich et al. 2016).

### Whole-genome sequencing breakpoint analysis and G4 overlap

Mutation breakpoints were determined using the start and end sites of SVs and from start sites of small indels. CNV regions were excluded from breakpoint analysis because breakpoint ends are not exactly determined in CNVkit but estimated from region coverage differences. Each mutation endpoint was adjusted to a region of 100bp or 1Kb centered on the endpoint in GenomicRanges (Lawrence et al. 2013). Experimental sequenced LiK and PDS G4 datasets were accessed from (Marsico et al. 2019) and computation PQS G4s were determined using sequences derived from the BSgenome.Hsapiens.UCSC.hg19 (v1.4.3) and Bsgenome.Mmusculus.UCSC.mm10 (v.1.4.0) packages and entered into pqsfinder (v2.4.0) (Hon et al. 2017) with minimum score of 20 used to identify all G4s in a given sequence region. Overlap analysis was performed in GenomicRanges. Visualizations were performed in GViz (v1.36.2) (Hahne F 2016). Overlap significance testing was performed using the regioneR package (v1.2.4) (Gel et al. 2016), to do permutation testing with 1000 iterations of random shuffling.

### Quantification and statistical analysis

ANOVA and t-tests for statistical significance was performed in Graphpad Prism software. Image quantification was performed as above in FIJI or Imaris and exported to Prism. Significance testing for G4 overlap was performed as above in regioneR (Gel et al. 2016). Descriptions of statistical analysis presented in the figures are within corresponding figure legends.

## Acknowledgements

We are grateful to all the members of the Smogorzewska and Hatten laboratories for helpful discussions throughout the project. We thank Alana C. Bernys, Madison Rex, Ksenia Morozova, Isaac Nathoo, Yakshi Dabas, and Rio McLellan for technical assistance on this project. We would like to thank Dr. Kärt Mätlik for bioinformatics analysis advice and manuscript comments. We thank Dr. Nathaniel Heintz for experimental advice on isolating RNA from fixed cerebellar nuclei sorted for EdU. We thank Dr. Maria Jasin for providing us with the *Brca2^Δex3-4^* mice. We gratefully acknowledge the Bioinformatics Resource Center, the Bio-Imaging Resource Center, the Flow Cytometry Resource Center, Genomics Resource Center, and the Comparative Bioscience Center at the Rockefeller University for experimental support.

This work was supported by the Emerald Foundation Research Grant (to A.S. and M.E.H.), Robertson Therapeutic Development Fund (to A.S. and D.L.K) and NIH/NCI Grant R01CA204127 (to A.S.), and National Institute of Neurological Disorders and Stroke (NINDS) 5R01NS116089 (to M.E.H.). Human MB work was supported by National Center for Advancing Translational Sciences UL1TR001866 (A.S.). D.L.K. was supported by the Rockefeller Women & Science Graduate Fellowship, Anderson Center Graduate Fellowship, and NIH Cancer Biology Training Grant CA009673. S.C.C. acknowledges support from the Intramural Research Program of the NIH National Human Genome Research Institute.

## Author Contributions

M.E.H. and A.S. conceptualized the study. A.S., M.E.H., and D.L.K. designed the experiments. D.L.K. performed all molecular experiments and isolated nucleic acids for next-generation sequencing. M.R.P. performed RNAseq alignment and differential gene expression analysis. M.A.S. did sequence alignment, variant calling, and filtering of WGS data. D.L.K., M.R.P., T.S.C., and A.L.H.W. performed further bioinformatics analysis. Y.F. isolated GCPs from animals at P7. T.F.W. performed mouse breeding and genotyping. S.S. assisted in EdU-positive tumor cell flow cytometry. S.C.C. supported WGS sequencing and analysis of tumor DNA. Clinical information about the FA-D1 patient from whom the HMB1 cells were generated was provided by M.L.M. and J.E.W. C.S.G. performed MB resection and coordinated with A.S. for tissue collection. D.L.K. and A.S. assembled the figures. D.L.K. and A.S. wrote the manuscript with input from all authors.

